# Multifaceted roles of SAMHD1 in cancer

**DOI:** 10.1101/2021.07.03.451003

**Authors:** Katie-May McLaughlin, Jindrich Cinatl, Mark N. Wass, Martin Michaelis

## Abstract

SAMHD1 is discussed as a tumour suppressor protein, but its potential role in cancer has only been investigated in very few cancer types. Here, we performed a systematic analysis of the TCGA (adult cancer) and TARGET (paediatric cancer) databases, the results of which did not suggest that SAMHD1 should be regarded as a *bona fide* tumour suppressor. SAMHD1 mutations that interfere with SAMHD1 function were not associated with poor outcome, which would be expected for a tumour suppressor. High SAMHD1 tumour levels were associated with increased survival in some cancer entities and reduced survival in others. Moreover, the data suggested differences in the role of SAMHD1 between males and females and between different races. Often, there was no significant relationship between SAMHD1 levels and cancer outcome. Taken together, our results indicate that SAMHD1 may exert pro-or anti-tumourigenic effects and that SAMHD1 is involved in the oncogenic process in a minority of cancer cases. These findings seem to be in disaccord with a perception and narrative forming in the field suggesting that SAMHD1 is a tumour suppressor. A systematic literature review confirmed that most of the available scientific articles focus on a potential role of SAMHD1 as a tumour suppressor. The reasons for this remain unclear but may include confirmation bias and publication bias. Our findings emphasise that hypotheses, perceptions, and assumptions need to be continuously challenged by using all available data and evidence.

## Introduction

Sterile α motif and histidine-aspartic domain containing protein 1 (SAMHD1) was initially discovered in dendritic cells and named dendritic cell-derived IFN-y induced protein (DCIP) [Li et al., 2000; Coggins et al., 2020]. SAMHD1 is indeed involved in the regulation of interferon signalling, and SAMHD1 mutations are associated with Aicardi-Goutieres syndrome, an autoimmune inflammatory disorder characterised by a dysregulated interferon response [Rice et al., 2009; Mauney & Hollis, 2018; Coggins et al., 2020].

In the meantime, SAMHD1 has been shown to exert a range of additional functions [Mauney & Hollis, 2018; Coggins et al., 2020; Chen et al., 2021]. As a deoxynucleoside triphosphate hydrolase (dNTPase), that cleaves deoxynucleoside triphosphates (dNTPs) into deoxynucleosides and triphosphate, SAMHD1 plays, together with enzymes that catalyse dNTP biosynthesis, an important role in the maintenance of balanced cellular dNTP pools [Mauney & Hollis, 2018; Coggins et al., 2020; Chen et al., 2021]. Since imbalances in cellular dNTP pools affect cell cycle regulation and DNA stability, SAMHD1 is also involved in the regulation of these processes [Chen et al., 2021].

In addition to controlling cellular dNTP levels, SAMHD1 has been shown to maintain genome integrity by a range of further mechanisms, including maintenance of telomere integrity, inhibition of LINE-1 retrotransposons, facilitation of homologous recombination-mediated double-strand break repair and DNA end joining, and prevention of R-loop formation at transcription-replication conflict regions [Herold et al., 2017a; Akimova et al., 2021; Chen et al., 2021; Park et al., 2021]. Additionally, low SAMHD1 levels have been detected in chronic lymphocytic leukaemia (CLL), lung cancer, cutaneous T-cell lymphoma, AML, colorectal cancer, and Hodgkin lymphoma. Moreover, loss-of-function SAMHD1 mutations have been described in cancer types including CLL and colorectal cancer [Herold et al., 2017a; Mauney & Hollis, 2018; Coggins et al., 2020; Chen et al., 2021]. Due to these observations, SAMHD1 is being considered as a tumour suppressor protein.

However, the potential role of SAMHD1 in cancer diseases is more complex. It also recognises and cleaves the triphosphorylated, active forms of a range of anti-cancer nucleoside analogues. In this context, SAMHD1 has been described as a clinically relevant resistance factor in acute myeloid leukaemia (AML) and acute lymphoblastic leukaemia (ALL) against nucleoside analogues including cytarabine, decitabine, and nelarabine [Herold et al., 2017b; Schneider et al., 2017; Knecht et al., 2018; Oellerich et al., 2019; Rothenburger et al., 2020].

So far, the potential tumour suppressor activity of SAMHD1 has only been investigated in a few cancer types. To establish a broader understanding of the role of SAMHD1 in cancer, we here performed a systematic analysis of mutation data, gene expression data, and cancer patient survival data provided by The Cancer Genome Atlas (TCGA) [Cancer Genome Atlas Research Network, 2008] and the Therapeutically Applicable Research To Generate Effective Treatments (TARGET) (https://ocg.cancer.gov/programs/target) databases. The TCGA provided data from 9,703 patients with 33 different types of adult cancer and the TARGET database from 1,091 patients with seven different paediatric cancer types.

## Results

### High *SAMHD1* expression is not consistently associated with increased survival

Although SAMHD1 has recently been considered as a tumour suppressor protein [Herold et al., 2017a; Mauney & Hollis, 2018; Coggins et al., 2020], high *SAMHD1* expression in tumour tissues was not associated with favourable outcomes across all patients in the TCGA (Figure 1A). In contrast, high *SAMHD1* expression was associated with favourable outcome across all patients in the paediatric cancer database TARGET (Figure 1A).

**Figure 1.**
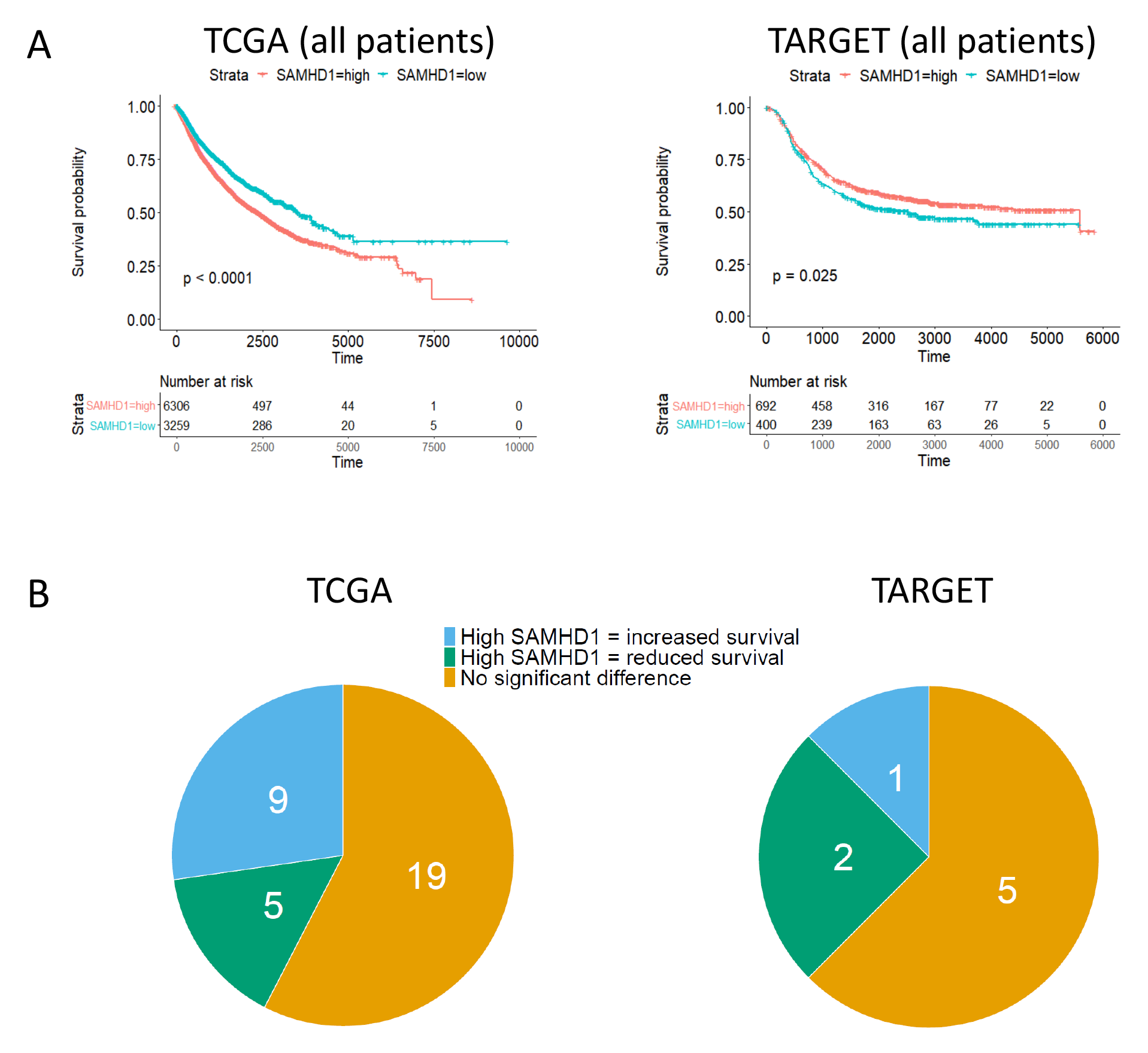
Effect of *SAMHD1* expression in cancer patients. A) Kaplan Meier plots indicating survival in cancer patients with tumours characterised by high or low *SAMHD1* expression (as determined by best separation) across all patients in the TCGA and TARGET databases. P-values were determined by log-rank test. B) Pie charts indicating the number of cancer types for which high SAMHD1 expression was associated with increased survival, reduced survival, or not significantly associated with survival based on data from the TCGA and TARGET databases. Data are presented in Supplementary Table 1 and Supplementary Table 2.

Considering the individual cancer categories in the TCGA database, high *SAMHD1* expression was significantly associated with increased survival in nine out 33 of cancer categories (Figure 1B, Supplementary Table 1) and with poor outcome in five cancer categories (Figure 1B, Supplementary Table 1). This indicates that the role of SAMHD1 differs between cancer types and that it does not always function as a tumour suppressor. Similarly, high SAMHD1 expression was significantly correlated with longer survival in only one cancer type (osteosarcoma) in the TARGET database but with reduced survival in two others (acute lymphoblastic leukaemia, Wilm’s tumour) (Figure 1B, Supplementary Table 2).

### Role of *SAMHD1* expression in the context of sex

Next, we analysed the role of SAMHD1 in males and females in TCGA and TARGET. In TCGA, there were no sex-specific differences with regard to the association of SAMHD1 with survival time across all cancer types (Figure 2A). However, some discrepancies became visible upon the comparison of the role of SAMHD1 in the 27 cancer entities that occur in both females and males (Figure 2B, Supplementary Table 3). When we did not consider statistical significance levels, high SAMHD1 levels were associated with higher 5-year survival rates in 13 cancer entities across all patients, in 17 cancer entities in female patients, and in 14 cancer entities in male patients (Figure 2B, Supplementary Table 3).

**Figure 2.**
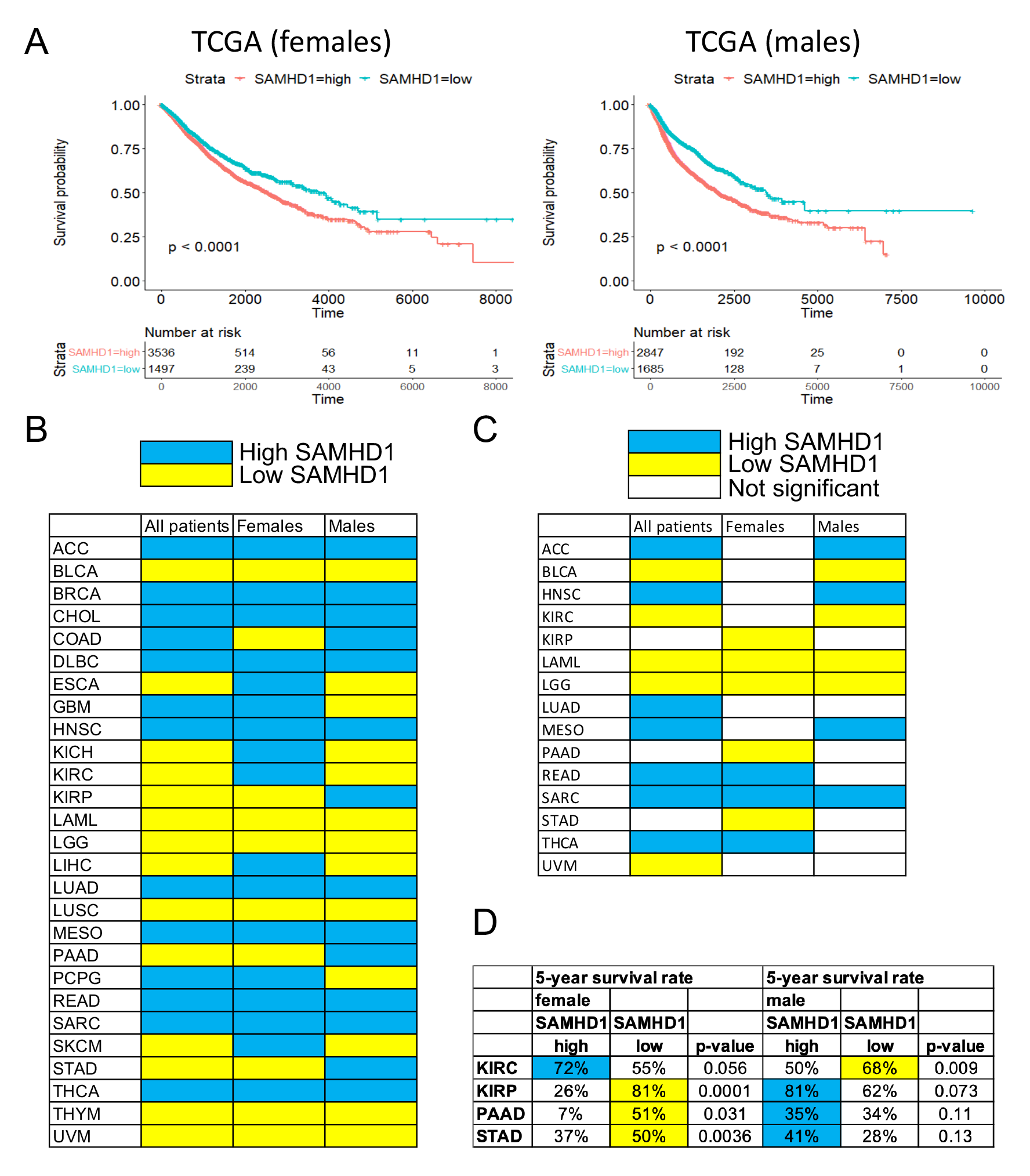
*SAMHD1* expression levels and 5-year survival rates in dependence of sex based on TCGA data. A) Kaplan Meier plots indicating sex-specific survival in cancer patients with tumours characterised by high or low *SAMHD1* expression (as determined by best separation). P-values were determined by log-rank test. B) Heatmap indicating the association of *SAMHD1* expression and 5-year survival rates (blue: high SAMHD1 associated with higher survival rates, yellow: low SAMHD1 associated with higher survival rates). C) Heatmap indicating cancer entities in which high *SAMHD1* expression (blue) or low *SAMHD1* expression (yellow) is significantly (p<0.05) associated with higher 5-year survival rates. D) Cancer entities in which SAMHD1 displays a trend towards differing roles by sex. Blue indicates higher survival rates in patients with tumours with high SAMHD1 levels, yellow in patients with low SAMHD1 levels. Abbreviations: ACC, Adrenocortical carcinoma; BLCA, Bladder Urothelial Carcinoma; BRCA, Breast invasive carcinoma; CESC, Cervical squamous cell carcinoma and endocervical adenocarcinoma; CHOL, Cholangiocarcinoma; COAD, Colon adenocarcinoma; DLBC, Lymphoid Neoplasm Diffuse Large B-cell Lymphoma; ESCA, Oesophageal carcinoma; GBM, Glioblastoma multiforme; HNSC, Head and Neck squamous cell carcinoma; KICH, Kidney Chromophobe; KIRC, Kidney renal clear cell carcinoma; KIRP, Kidney renal papillary cell carcinoma; LAML, Acute Myeloid Leukaemia; LGG, Low Grade Glioma; LIHC, Liver hepatocellular carcinoma; LUAD, Lung adenocarcinoma; LUSC, Lung squamous cell carcinoma; MESO, Mesothelioma; OV, Ovarian serous cystadenocarcinoma; PAAD, Pancreatic adenocarcinoma; PCPG, Pheochromocytoma and Paraganglioma; PRAD, Prostate adenocarcinoma; READ, Rectum adenocarcinoma; SARC, Sarcoma; SKCM, Skin Cutaneous Melanoma; STAD, Stomach adenocarcinoma; TGCT, Testicular Germ Cell Tumours; THCA, Thyroid carcinoma; THYM, Thymoma; UCEC, Uterine Corpus Endometrial Carcinoma; UCS, Uterine Carcinosarcoma; UVM, Uveal Melanoma.

When we only considered comparisons in which the 5-year survival rates were significantly different (p<0.05) between high and low SAMHD1-expressing tumours for at least one comparison (across all patients, in females, and/ or males), differences reached significance for only one sex in ten cancer types (Figure 2C, Supplementary Table 3). Consistent findings were obtained for three cancer types (LAML, LGG, SARC, all abbreviations for cancer entities are provided in Supplementary Table 1 and the legend of Figure 2).

Although differences did not reach our cut-off value for statistical significance (p < 0.05), trends were detected indicating opposite effects of SAMHD1 on disease outcome between the sexes in four cancer entities, (Figure 2D, Supplementary Table 3). In kidney renal clear cell carcinoma (KIRC), low SAMHD1 levels were associated with a higher 5-year survival rate (68%) than high SAMHD1 levels (50%) in males (p=0.009). In contrast, high SAMHD1 levels were related to higher survival in females (72% vs. 55%), with the p-value being close to significance (p=0.056). In three other cancer types (KIRP, PAAD, STAD), low SAMHD1 levels were associated with higher 5-year survival in females and with lower 5-year survival in males (Figure 2D, Supplementary Table 3).

In TARGET, high SAMHD1 levels were significantly associated with increased survival across all cancer types in females (Figure 3A) but not in males (Figure 3B). For seven paediatric cancer types, data were available for both sexes. When we did not consider statistical significance levels, higher 5-year survival rates were recorded for SAMHD1 patients with SAMHD1 high tumours across all patients, in one entity in female patients, and in four entities in male patients (Figure 3B, Supplementary Table 4.

**Figure 3.**
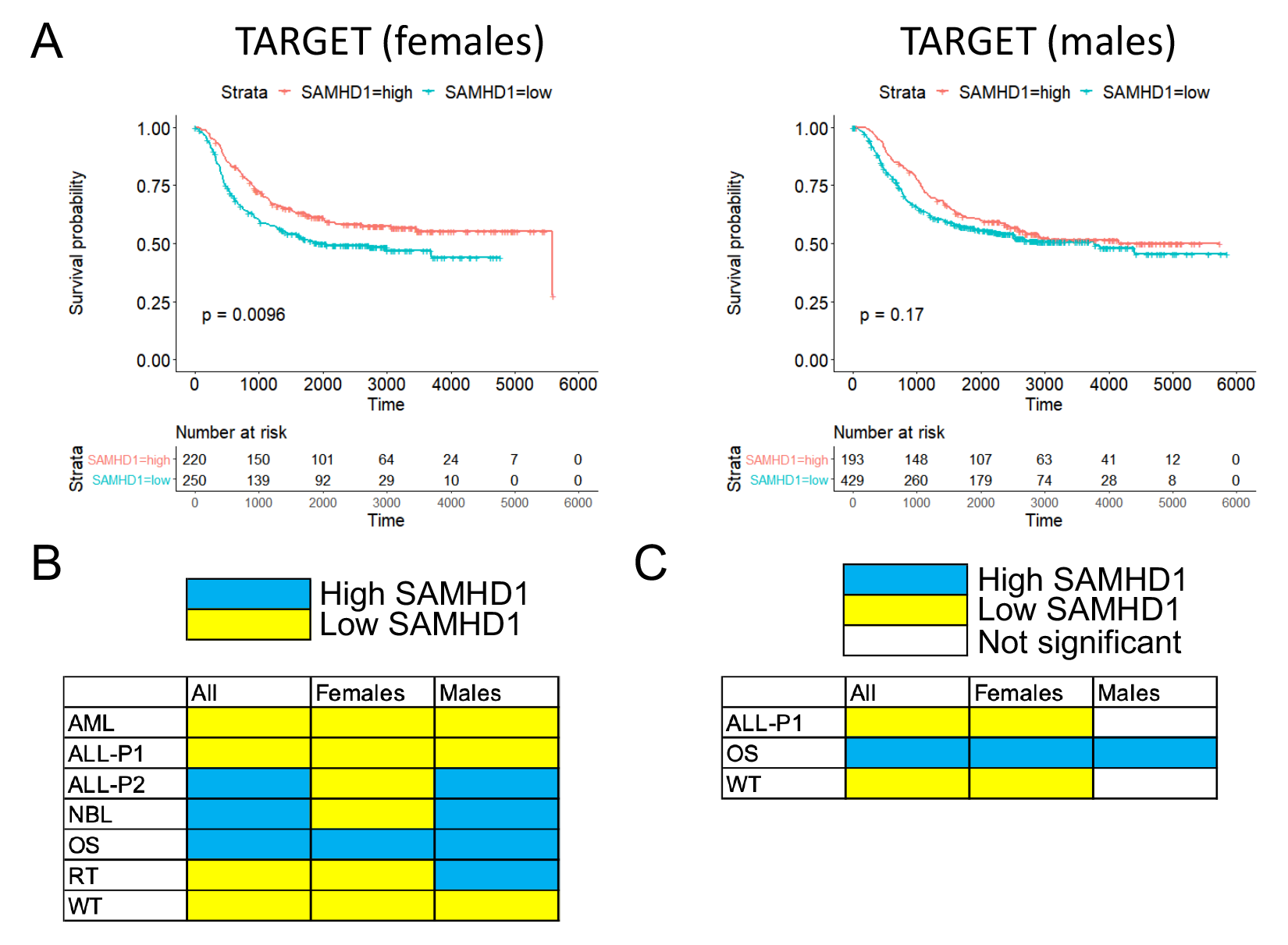
*SAMHD1* expression levels and 5-year survival rates in dependence of sex based on TARGET data. A) Kaplan Meier plots indicating sex-specific survival in cancer patients with tumours characterised by high or low *SAMHD1* expression (as determined by best separation). P-values were determined by log-rank test. B) Heatmap indicating the association of *SAMHD1* expression and 5-year survival rates (blue: high SAMHD1 associated with higher survival rates, yellow: low SAMHD1 associated with higher survival rates). C) Heatmap indicating cancer entities in which high *SAMHD1* expression (blue) or low *SAMHD1* expression (yellow) is significantly (p<0.05) associated with higher 5-year survival rates.

When we only considered cancer types in which the 5-year survival rates were significantly different (p<0.05) between high and low SAMHD1-expressing tumours for at least one comparison (across all patients, in females, and/ or males), differences reached significance only for females in two cancer types (Figure 3C, Supplementary Table 4). Taken together, these data suggest that the role of SAMHD1 in cancer may differ between the sexes in some cancer types.

Notably, the potentially different roles between SAMHD1 in female and male cancer patients do not appear to be the consequence of sex-specific discrepancies in *SAMHD1* expression. In TCGA, there were no significant sex-specific differences in *SAMHD1* expression between tumour samples and matched normal tissue samples from females and males, when we excluded sex-specific cancer types (CESC, OV, PRAD, TGCT, UCEC, UCS) and BRCA (only 12 out of 1,089 tumour tissue samples from male patients, only one out of 113 matched normal tissue samples from a male) (Supplementary Figure 1). SAMHD1 expression was also not significantly different in males and females in the TARGET database (Supplementary Figure 2).

### Role of *SAMHD1* expression in the context of race

The vast majority of data in TCGA and TARGET are derived from white individuals, which reduces the significance of the race-related data. In TCGA, high SAMHD1 expression was associated with reduced overall survival in white patients (Figure 4). This reflects the findings obtained across all patients (Figure 1A) and probably that 6,834 (82%) out of 8,319 patients, for whom race data are available, are reported to be white. Apart from this, a significant difference in outcome in dependence of tumour SAMHD1 levels was only detected in Native Hawaiian or other Pacific islander patients, in whom high SAMHD1 was associated with improved survival (Figure 4). However, only 13 individuals fell into this category. Cancer-type specific comparisons did not reveal significant differences in SAMHD1-related outcomes between racial groups (Supplementary Figure 3, Supplementary Table 5), which may be due to the low numbers of patients in most of the categories (Figure 4). SAMHD1 levels were generally similar between the different race groups (Supplementary Figure 4). Only Native Hawaiian or other Pacific islander patients displayed increased levels (Supplementary Figure 4).

**Figure 4.**
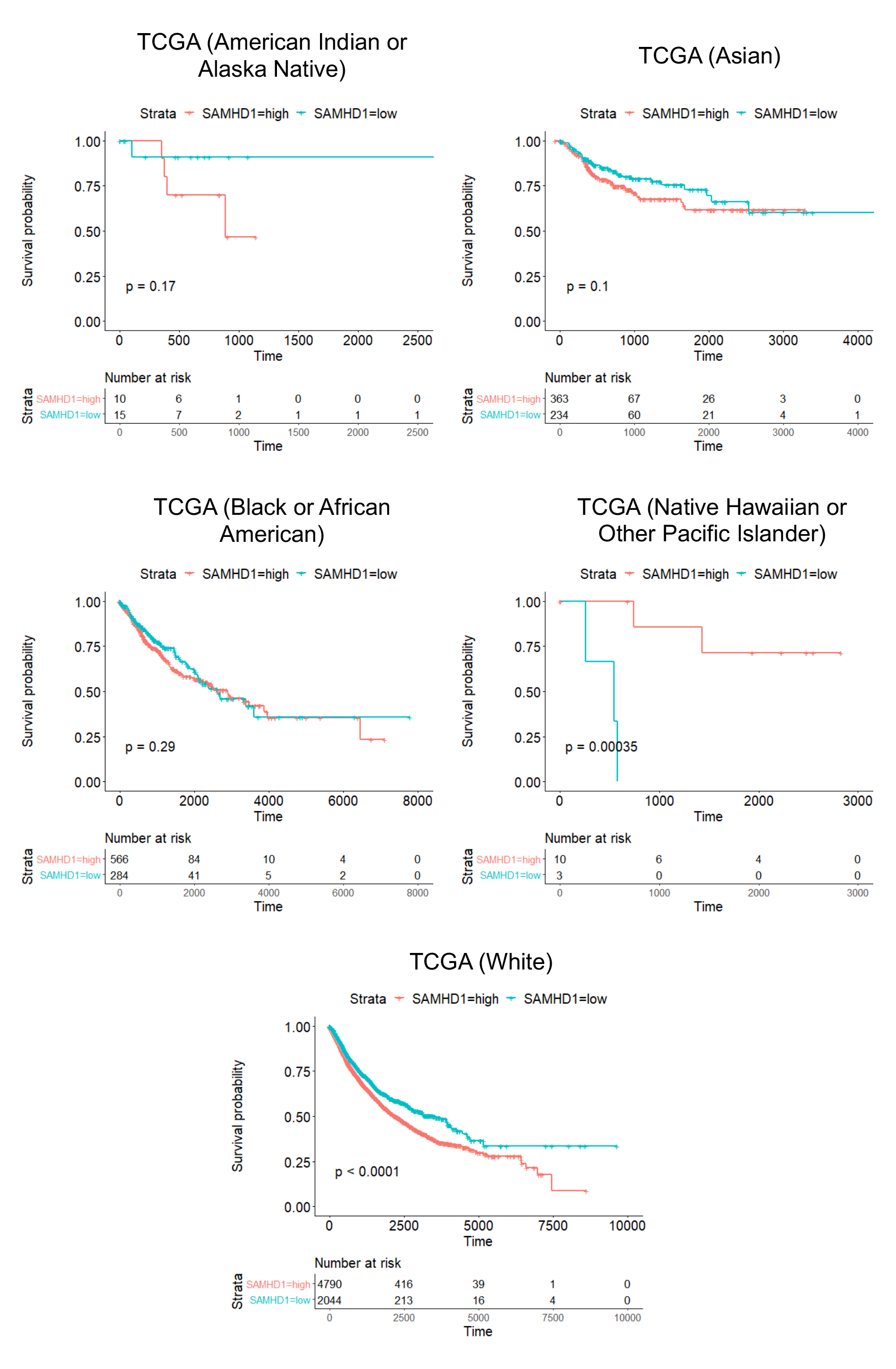
Kaplan-Meier plots indicating survival in cancer patients of different race with tumours characterised by high or low *SAMHD1* expression (as determined by best separation) based on TCGA data. P-values were determined by log-rank test.

Stratifying of patients in the TARGET database according to race provided some trends, which may point towards differences, but the numbers are too low to draw firm conclusions (Figure 5, Supplementary Figure 5, Supplementary Table 6). No significant differences were detected between the SAMHD1 levels in the different race groups (Supplementary Figure 6).

**Figure 5.**
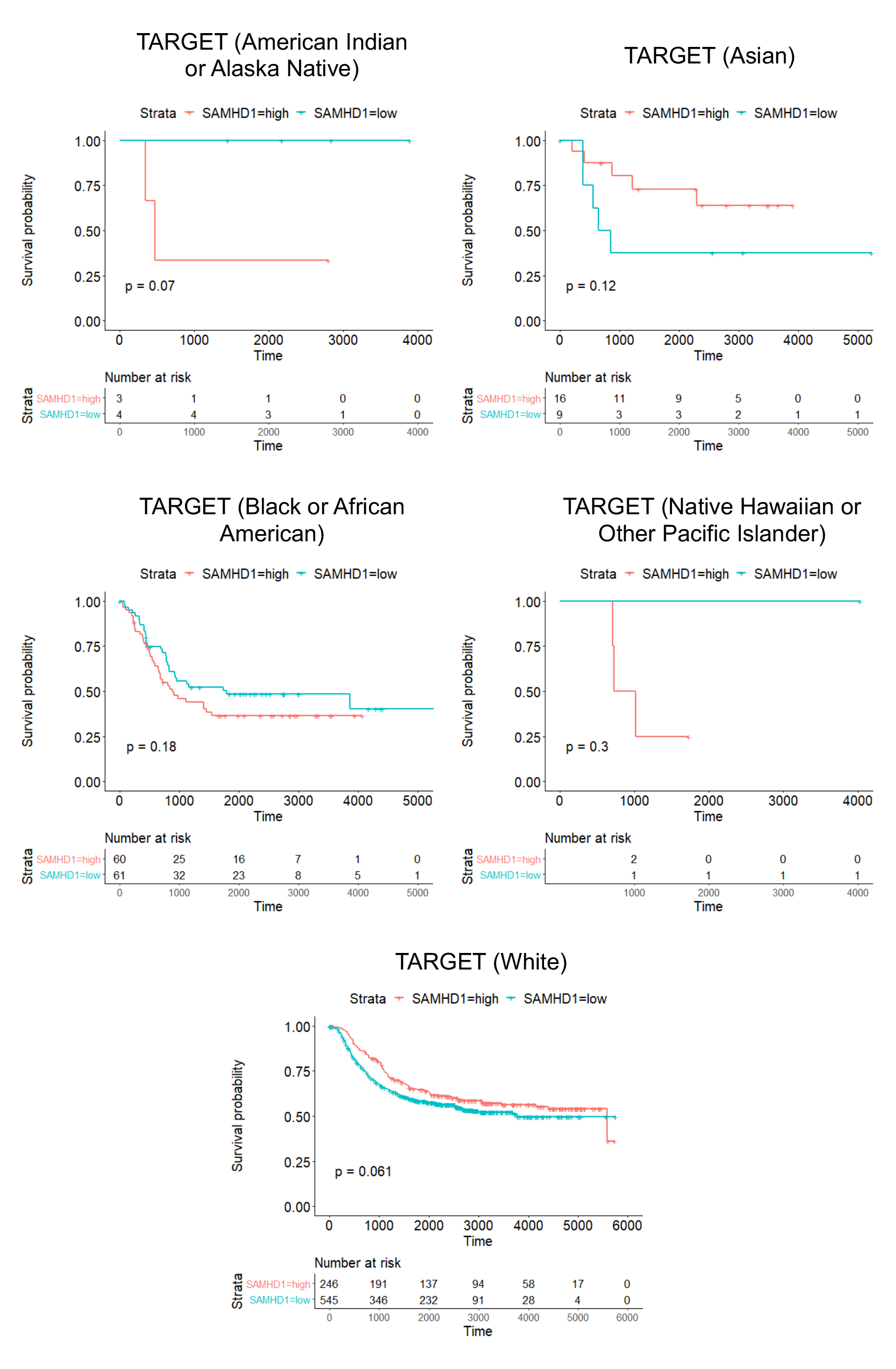
Kaplan-Meier plots indicating survival in cancer patients of different race with tumours characterised by high or low *SAMHD1* expression (as determined by best separation) based on TARGET data. P-values were determined by log-rank test.

### *SAMHD1* expression in tumour vs matched normal samples

To further investigate the potential role of SAMHD1 in cancer, we next compared *SAMHD1* expression data in tumour tissue and matched normal samples, which were available for 695 patients and 21 cancer types in TCGA. Across all patients, there was no significant difference between the *SAMHD1* FPKM (fragments per kilobase of transcript per million mapped reads) values of tumour samples and matched normal samples (Wilcoxon signed-rank test p-value = 0.14). However, when stratifying by cancer type, SAMHD1 levels significantly differed (p<0.05) between tumour samples and matched control samples in seven cancer types (Supplementary Table 7). SAMHD1 was higher in matched control samples suggesting tumour suppressor activity in three cancer types (BLCA, LUAD, LUSC) and higher in tumour samples from four cancer types (KIRC, KIRP, LIHC, STAD) suggesting oncogenic action (Supplementary Table 7).

Next, we compared the results on potential tumour suppressor or oncogenic functions of SAMHD1 from tumour and matched normal tissues (Supplementary Table 7) to those obtained from analysing 5-year survival in cancer patients with SAMHD1 low or high tumours (Figure 2; Supplementary Table 1). When we did not consider statistical significance levels, SAMHD1 levels were higher in control tissues suggesting tumour suppressor activity in ten cancer entities (Figure 6A, Supplementary Table 7). In twelve of the 21 cancer types, both SAMHD1 levels in tumour and matched normal tissues and the relationship of 5-year survival and tumour SAMHD1 levels indicated a similar role of SAMHD1, i.e. tumour suppressor (higher *SAMHD1* expression in matched normal tissue, higher 5-year survival in patients with SAMHD1 high tumours) or oncogenic (higher *SAMHD1* expression in tumour tissues, higher 5-year survival in patients with SAMHD1 low tumours) activity (Figure 6A, Supplementary Table 7).

**Figure 6.**
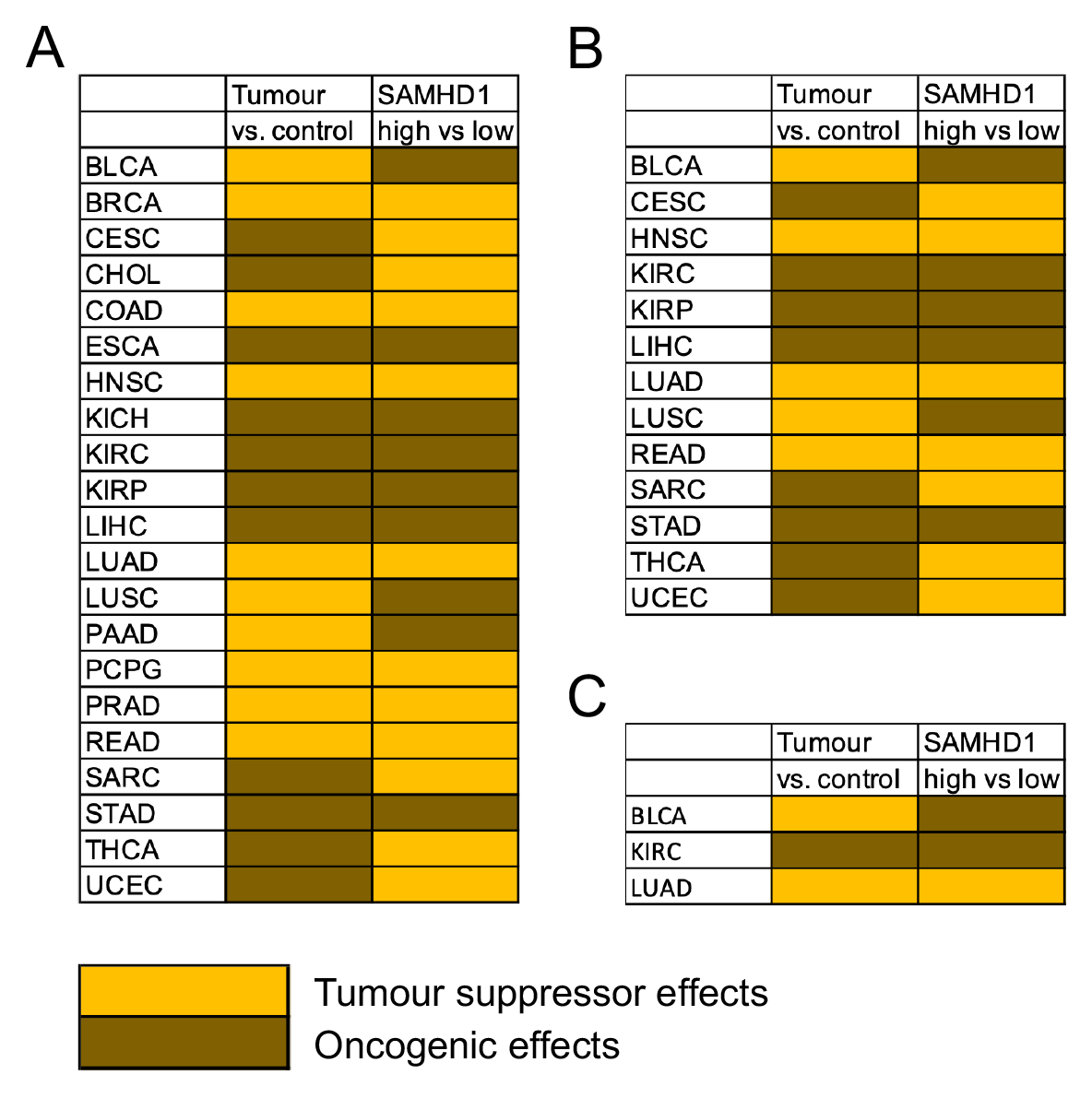
Tumour suppressor and oncogenic effects of SAMHD1 in different cancer types, as suggested by SAMHD1 levels in tumour tissues vs. matched normal tissues (Tumour vs. control) or the comparison of 5-year survival in patients with SAMHD1 high or low tumours (SAMHD1 high vs. low). Higher SAMHD1 levels in matched normal tissues were interpreted as tumour suppressor activity, while higher SAMHD1 levels in tumour tissues as indication of oncogenic effects. Higher 5-year survival in patients with SAMHD1 high tumours was construed as sign of tumour suppressor activity, higher 5- year survival in patients with SAMHD1 low tumours indication of oncogenic effects. A) Data for all available comparisons. B) Data for entities, in which at least the difference for one comparison reached statistical significance. C) Data for entities, in which the difference for both comparisons reached statistical significance.

In the next step, only cancer entities were considered for which at least one of the comparisons had resulted in a statistically significant (p<0.05) difference, leaving 13 cancer types (Figure 6B, Supplementary Table 7). In seven of these 13 cancer types, the anticipated role of SAMHD1 (tumour suppressor or oncogenic) coincided between both comparisons (Figure 6B, Supplementary Table 7).

In only three cancer entities (BLCA, KIRC, LUAD), the differences reached statistical significance for both comparisons (Figure 6C, Supplementary Table 7). SAMHD1 consistently displayed oncogenic activity in KIRC (Kidney renal clear cell carcinoma) and tumour suppressor activity in LUAD (Lung adenocarcinoma). In BLCA (Bladder Urothelial Carcinoma), higher SAMD1 levels in matched normal tissue samples suggested tumour suppressor activity, whereas higher 5-year survival in patients with SAMHD1 low tumours suggested oncogenic effects (Figure 6C, Supplementary Table 7). Hence, SAMHD1 may exert oncogenic activity in KIRC and tumour suppressor activity in LUAD, but clear evidence is lacking for other cancer entities.

### SAMHD1 regulation by methylation and miRNAs

Promotor methylation and miRNAs have been described to be involved in SAMHD1 regulation [de Silva et al., 2013; Kohnken et al., 2017; Chen et al., 2021]. Tumour and normal sample SAMHD1 expression and promoter methylation beta values were available for 18 cancer types in TCGA. SAMHD1 promotor methylation significantly inversely correlated with SAMHD1 expression levels across all patients, but the correlation coefficient was moderate and the relationship appears weak (Figure 7A).

**Figure 7.**
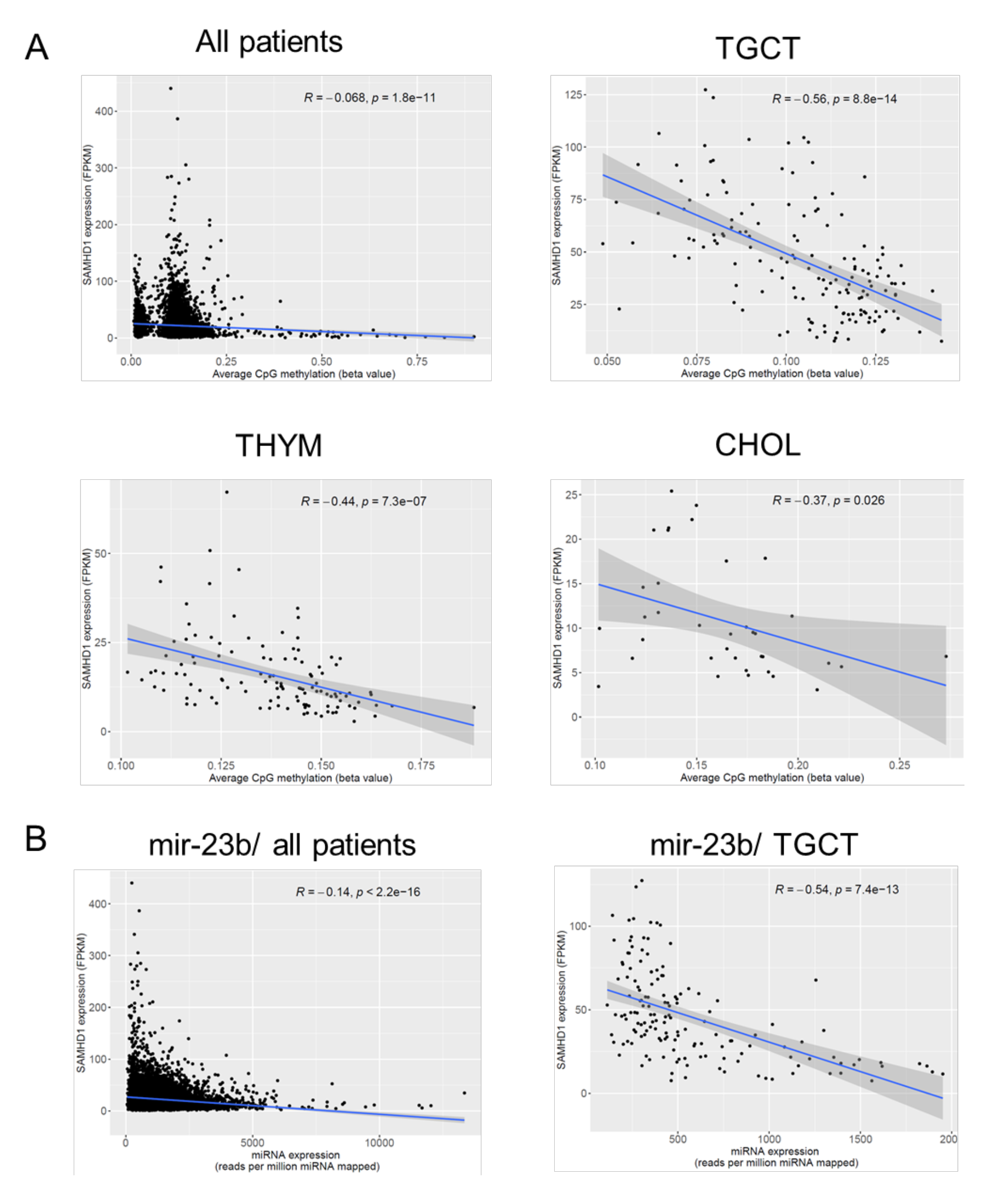
Inverse correlation between SAMHD1 promotor methylation levels or miRNA levels and SAMHD1 expression based on TCGA data. A) Correlation between SAMHD1 promotor methylation levels and SAMHD1 expression across all patients and in THYM patients, which displayed the strongest inverse correlation across all cancer types. Data for all cancer types are presented in Supplementary Table 8. B) Correlation of mir-23b with SAMHD1 expression across all patients and of mir-30c-1 with SAMHD1 in THYM. mir-23b was the miRNA that displayed the strongest inverse correlation with SAMHD1 across all patients. The inverse correlation between mir-30c-1 and SAMHD1 was the strongest among all miRNAs in all cancer types. Data for all significant inverse correlations of miRNAs and SAMHD1 across all cancer types are provided in Supplementary Table 10.

When we looked at the individual cancer types, an inverse correlation between SAMHD1 expression and promotor methylation was detected in 18 cancer entities (Supplementary Table 8). In eleven of these cancer types, the inverse correlations displayed p-values < 0.05 (Supplementary Table 8). The cancer types with the strongest inverse correlations between SAMHD1 expression and promotor methylation were TGCT (Testicular Germ Cell Tumours), THYM (Thymoma), and CHOL (Cholangiocarcinoma) (Figure 7A). There were also 15 cancer entities with a direct correlation between SAMHD1 expression and promotor methylation, only four of which were associated with a p-value < 0.05. These data suggest that promotor methylation is one SAMHD1 regulation mechanism among others and that its role differs between cancer types.

Eight miRNAs (mir-30a, mir-155, and six subtypes of mir-181) have been described to be involved in SAMHD1 regulation [Jin et al., 2014; Pilakka-Kanthikeel et al., 2015; Kohnken et al., 2017; Riess et al., 2017]. Moreover, 21 miRNAs were indicated to interact with SAMHD1 in the DIANA-TarBase v8 [http://www.microrna.gr/tarbase] [Karagkouni et al., 2018], an online resource that lists experimentally validated miRNA/mRNA interactions. After the removal of overlaps, this resulted in a list of 28 miRNAs with a documented effect on SAMHD1 (Supplementary Table 9).

Of these 28 miRNAs, 26 miRNAs were found to be inversely correlated with SAMHD1 expression in one (mir-21) to 18 (mir-183) cancer entities (Supplementary Table 10). Six miRNAs (mir-23b, mir-30a, mir-192, mir-181d, mir-218-1, mir-218-2) were significantly (p<0.05) inversely correlated with SAMHD1 across all patients, with mir-23b showing the strongest inverse correlation (R= −0.14, p=2.2×10^-16^) (Figure 7B, Supplementary Table 10). The strongest inverse correlation was detected between mir-30c-1 and SAMHD1 in THYM (R= −0.85, p=0.02) (Figure 7B, Supplementary Table 10).

Each of these 28 miRNAs were found to be inversely correlated with SAMHD1 expression in between two (mir-155) and all 28 (mir-23b and mir-183) cancer entities (Supplementary Table 10). Six miRNAs (mir-23b, mir-30a, mir-192, mir-181d, mir-218-1, mir-218-2) were significantly (p<0.05) inversely correlated with SAMHD1 across all patients, with mir-23b showing the strongest inverse correlation (Figure 7B, Supplementary Table 10). The strongest inverse correlation in a cancer type was detected between mir-23b and SAMHD1 in TGCT (R= −0.54, p=7.37×10^-13^) (Figure 7B, Supplementary Table 10).

Taken together, SAMHD1 levels are determined by complex regulation mechanisms that include promotor methylation and miRNAs, together with post-translational modifications such as phosphorylation and acetylation that have also been described [Coggins et al., 2020; Chen et al., 2021].

### *SAMHD1* mutations and patient survival

*SAMHD1* mutations have been described in cancers including chronic lymphocytic leukaemia, T-cell prolymphocytic leukaemia, mantle cell lymphoma, cutaneous T-cell lymphoma and colon cancer [Clifford et al., 2014; Guièze et al., 2015; Merati et al., 2015; Amin et al., 2016; Rentoft et al., 2016; Burns et al., 2018; Johansson et al., 2018; Nadeu et al., 2020; Bühler et al., 2021; Roider et al., 2021].

Mutation data was available for 10,149 patients in the TCGA. 15,351 out of 21,156 genes harboured at least one non-synonymous mutation in one patient (Supplementary Table 11). The three most commonly mutated genes were *TTN*, *MUC16*, and *TP53* (Supplementary Table 11). *TTN* and *MUC16* encode the two longest human proteins (36,800 and 14,500 amino acids, respectively) that are frequently found mutated. Mutations in these genes are commonly regarded not to be of functional relevance and removed as artefacts or used as indicators of the mutational burden of tumours, while *TP53* is known to be the most commonly mutated tumour suppressor gene [Lawrence et al., 2013; Kim et al., 2017; Levine, 2020; Oh et al., 2020; Wang et al., 2020; Yang et al., 2020].

In total, *SAMHD1* was mutated 201 times, including 175 non-synonymous mutations in 159 patients (1.57% of patients for whom mutation data was available) (Supplementary Table 11). This places *SAMHD1* within the top 15.3% of most commonly mutated genes (Figure 8A, Supplementary Table 11). Among the 135 patients with *SAMHD1* mutant tumours for whom survival data were available, *SAMHD1* mutations were associated with superior outcome (Figure 8B, Supplementary Table 12). In 18 of the 25 cancer types, in which *SAMHD1* mutations were detected, 5-year survival was higher in patients with *SAMHD1* mutant tumours (Supplementary Table 12). However, the significance of these data is limited due to the low number of SAMHD1 mutations. Notably, the p-value (0.07) was close to significance in UCEC, the cancer type with the most SAMHD1 mutations (35/ 6.6% out of 527), in which 93.2% of patients with SAMHD1 mutant cancers survived for five years, in contrast to 76.1% of the 492 UCEC patients with SAMHD1 wild-type cancers (Figure 8C, Supplementary Table 12).

**Figure 8.**
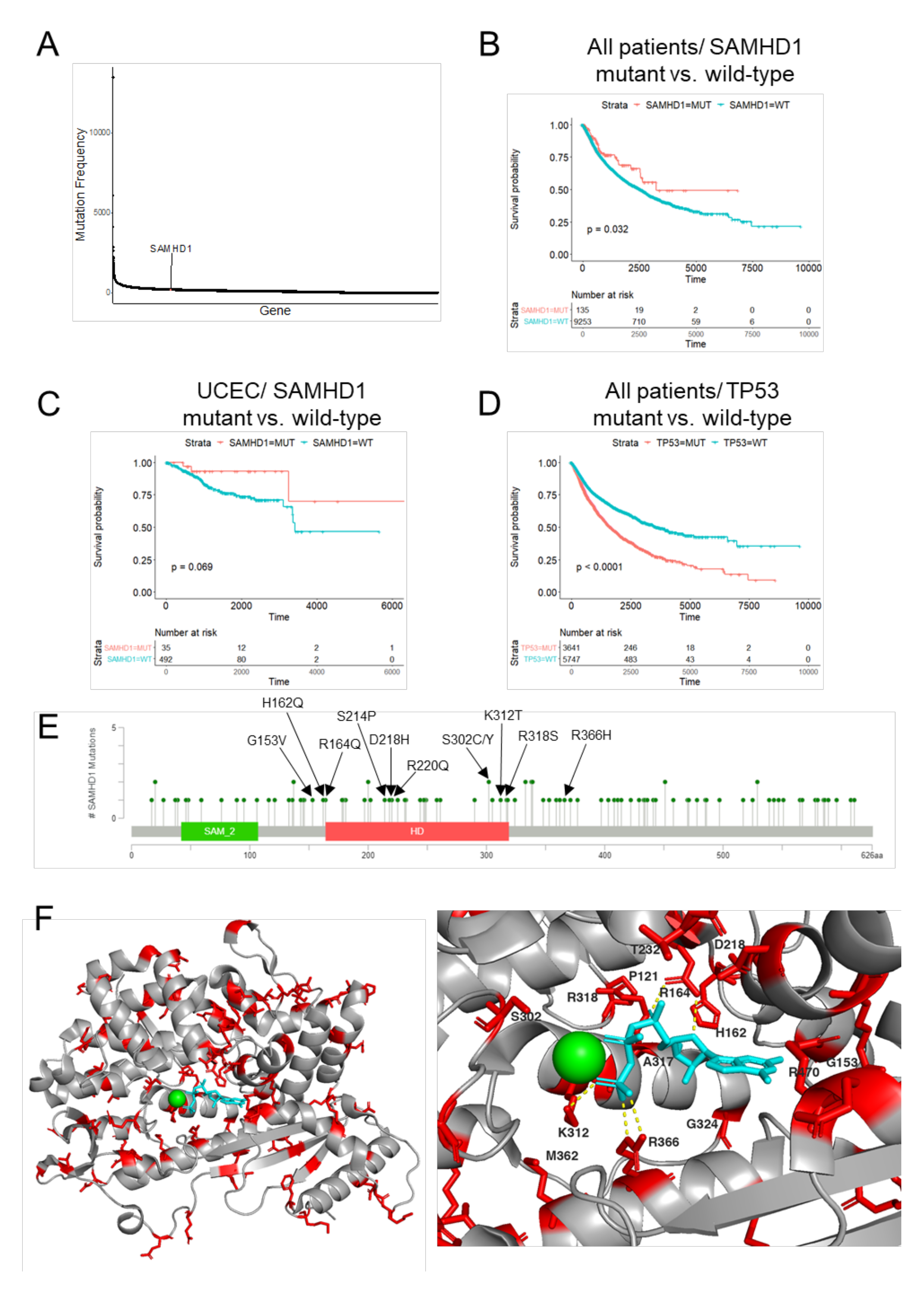
*SAMHD1* mutations in cancer tissues. A) *SAMHD1* was mutated 201 times, including 175 non-synonymous mutations, which puts *SAMHD1* within the 15.3% of most commonly mutated genes. B) Survival in patients with and without *SAMHD1* mutant tumours. C) Survival in UCEC (cancer type with the most *SAMHD1* mutations) patients with and without *SAMHD1* mutant tumours. D) Survival in patients with tumours with or without mutations in *TP53*, the most commonly mutated tumour suppressor genes. E) Lollipop plot indicating locations of missense mutations in *SAMHD1*. Residues predicted to be involved in ligand binding are labelled in bold. F) nonsynonymous mutations mapped (coloured red) onto the SAMHD1 protein structure (Protein Databank identifier 6DWD [Knecht et al., 2018] with bound clofarabine hydrochloride (indicated in cyan) and magnesium ion (green). The image on the left shows the full structure and on the right the active site is displayed. Yellow dashed lines indicate hydrogen bonds between mutated residues and ligand.

Although it is not possible to draw firm conclusions from these data, they do not support a general tumour suppressive role of SAMHD1, as mutations in tumour suppressor genes would rather be expected to result in shorter survival. For example, mutations in *TP53*, the most commonly mutated tumour suppressor gene [Levine, 2020], were associated with reduced survival (Figure 8D).

### *SAMHD1* mutations are likely to be deleterious

Twenty-nine of the mutations are likely to result in a loss of function, including 11 stop-gain, 11 frameshift, six splice site and one stop loss mutation. While 21 mutations were located in untranslated regions (six 5’ UTR, 15 3’ UTR), four were in introns, three in-frame, 25 were synonymous, with the remaining 104 resulting in nonsynonymous mutations.

50 mutations had already been described in cancer cells or were present in positions that had been found mutated in cancer cells (Supplementary Table 11). Three of the *SAMHD1* mutations identified in the TCGA (R143C/ UCEC patient, R145Q/ COAD patient, R290H/ STAD patient) were loss-of-function mutations associated with Aicardi-Goutiѐres syndrome [Rice et al., 2009; Mauney & Hollis, 2018; Coggins et al., 2020; UniProt Consortium, 2021]. 18 nonsynonymous mutations occurred at positions demonstrated to be important for SAMHD1 function by mutagenesis studies according to UniProt (Figure 8E, Figure 8F, Supplementary Table 11). This was supported by structural analysis which showed that ten non-synonymous mutations were located around the SAMHD1 active ligand binding sites (Figure 8F).

Next, the SAMHD1 non-synonymous variants were analysed using SIFT [Sim et al., 2012], PolyPhen-2 [Ng & Henikoff, 2001; Sim et al., 2012; Vaser et al., 2016], Condel [González-Pérez & López-Bigas, 2011], and CADD [Kircher et al., 2014] to predict if they are likely to have an effect on protein function (Supplementary Table 11). Approximately half of the amino acid changes were predicted to have a significant impact on SAMHD1 function (SIFT: 63/104 (60.6%), Polyphen-2: 50/104 (48.1%) and Condel: 54/104 (51.9%)). 72 of these variants also had a scaled CADD score of >20, which rates a variant among the top 1% of the most deleterious changes. Five variants displayed CADD scores >30 (Figure 8E). 39 variants had a SIFT rating of ‘tolerated’, a PolyPhen-2 rating of ‘benign’ and a Condel rating of ‘neutral’, of which 13 had a scaled CADD score of <10. Predictions for the remaining 17 amino acid changes were inconsistent (i.e. contrasting SIFT, PolyPhen-2 and Condel predictions) (Supplementary Table 11).

Taken together, many of the mutations appear to affect SAMHD1 function. However, loss-of-function should typically be associated with reduced survival in tumour suppressor genes.

### Literature review SAMHD1 and cancer

Our analysis of TCGA and TARGET data do not suggest that SAMHD1 generally functions as a tumour suppressor protein. Neither *SAMHD1* mutations nor low SAMHD1 levels were consistently associated with reduced survival. However, SAMHD1 is discussed as a potential tumour suppressor protein in the literature [Herold et al., 2017a; Chen et al., 2021]. Next, we performed a systematic review to compare our findings to those from the literature and to gain further insights into the narrative underlying the perceived role of SAMHD1 in cancer.

The literature search was performed in PubMed (https://pubmed.ncbi.nlm.nih.gov) on 17^th^ June 2021 using the search term "(((Cancer) OR (tumor) OR (tumour))) AND (SAMHD1)". It resulted in 150 hits, including 35 articles with relevant original data and 15 relevant secondary literature articles (reviews, editorials, comments) (Supplementary Figure 7, Supplementary Table 13).

The first articles reported on a potential role of SAMHD1 in cancer in 2013 [Clifford et al., 2014; de Silva et al., 2014; Shi et al., 2014]. The first one reported on low SAMHD1 levels in patients with Sézary syndrome, an aggressive subtype of cutaneous T-cell lymphoma, due to SAMHD1 methylation [de Silva et al., 2014], while the second paper described SAMHD1 variants associated with hepatitis B virus-and hepatitis C virus-induced hepatocellular carcinoma [Shi et al., 2014]. The third paper found SAMHD1 mutations in chronic lymphocytic leukaemia and proposed that these mutations promote leukaemia development by affecting SAMHD1-mediated DNA repair [Clifford et al., 2014].

Among the 35 articles that reported on an association between SAMHD1 and cancer, four did not (entirely) support the ‘SAMHD1 is a tumour suppressor’ narrative [Yang et al., 2017; Kodigepalli et al., 2018; Shang et al., 2018; Xagoraris et al., 2021]. One study correlated high *SAMHD1* levels with metastasis formation in colorectal cancer [Yang et al., 2017]. Kodigepalli et al., [2018] reported that SAMHD1 knock-out increases acute myeloid leukaemia cell proliferation via PI3K signalling but inhibits tumourigenesis potentially due to a lack of SAMHD1-mediated TNFalpha suppression. Notably, the title of this study exclusively focused on the inhibitory effects of SAMHD1 on leukaemia cell proliferation and did not mention its role in tumourigenesis [Kodigepalli et al., 2018]. One study detected a SAMHD1 increase upon lung cancer progression [Shang et al., 2018], and the most recent study, correlated the presence of SAMHD1 in Hodgkin lymphoma cells with unfavourable outcome [Xagoraris et al., 2021]. Notably, this study [Xagoraris et al., 2021] even referred to SAMHD1 as "novel tumour suppressor" in the title, although the study rather indicated an oncogenic role of SAMHD1.

The remaining articles largely focused on SAMHD1 mutations and reduced SAMHD1 levels in different cancer types as well as on SAMHD1’s potential role as a tumour suppressor involved in DNA repair (Supplementary Table 13). *SAMHD1* mutations were detected in patients with hepatocellular carcinoma [Shi et al., 2014], chronic lymphocytic leukaemia [Clifford et al., 2014; Guièze et al., 2015; Amin et al., 2016; Kim et al., 2016; Burns et al., 2018], cutaneous T-cell lymphoma [Merati et al., 2015], colorectal cancer [Rentoft et al., 2016], T-cell prolymphocytic leukaemia [Johansson et al., 2018], acute myeloid leukaemia [Zhu et al., 2018], and mantle cell lymphoma [Nadeu et al., 2020; Bühler et al., 2021].

In our TCGA analysis performed above (Figure 8, Supplementary Table 12), there is rather a trend towards higher 5-year survival rates among hepatocellular carcinoma (LIHC) patients with *SAMHD1* mutant tumours, although the significance of the data is limited due to low numbers (Supplementary Table 12). All four patients with *SAMHD1* mutant hepatocellular carcinoma survived for five years, while only 51.4% of 359 patients with *SAMHD1* wild-type hepatocellular carcinomas survived for five years. In colorectal adenocarcinoma (COAD), there was no noticeable difference between the survival of patients with *SAMHD1* mutant and *SAMHD1* wild-type tumours (Supplementary Table 12). Only in rectal adenocarcinoma (READ), a trend suggested that patients with *SAMHD1* mutant tumours may have a worse outcome. None out of five patients with *SAMHD1* mutant tumours survived for five years, while 54.9% of 130 patients with *SAMHD1* wild-type tumours did (Supplementary Table 12).

Acute myeloid leukaemia (LAML) was the only cancer type in which a significant difference was detected between patients with *SAMHD1* mutant and wild-type cancer cells. Patients with *SAMHD1* mutant leukaemia cells had a higher 5-year survival rate (Supplementary Table 8). However, this is most probably not due to general oncogenic activity, but because lack of SAMHD1 function results in a higher activity of nucleoside analogues including cytarabine and decitabine that are SAMHD1 substrates and commonly used for LAML treatment [Schneider et al., 2017; Oellerich et al., 2019].

The study on colorectal cancer [Rentoft et al., 2016] was the only one that had used TCGA data. However, it only used TCGA data to identify mutations, but did not compare survival in patients with and without *SAMHD1* mutations.

*SAMHD1* expression levels have been suggested to impact on cutaneous T-cell lymphoma [de Silva et al., 2014], lung cancer [Wang et al., 2014; Shang et al., 2018], colorectal cancer [Yang et al., 2017], and acute myeloid leukaemia [Jiang et al., 2020]. Low SAMHD1 levels were described in cutaneous T-cell lymphoma and acute myeloid leukaemia cells [de Silva et al., 2014; Jiang et al., 2020], supporting a potential role as a tumour suppressor. TCGA did not contain data on *SAMHD1* expression in cutaneous T-cell lymphoma or acute myeloid leukaemia cells relative to control cells.

In lung cancer, conflicting results were reported. One study found that SAMHD1 is down regulated in lung cancer by methylation and inhibits tumour cell proliferation [Wang et al., 2014]. The other study reported that SAMHD1 levels increase in the serum of lung cancer patients upon progression [Shang et al., 2018]. Our analysis of SAMHD1 data found significantly higher 5-year survival rates in patients with tumours displaying high *SAMHD1* expression levels (Supplementary Table 1) supporting the first study. Notably, elevated SAMHD1 in the serum of lung cancer patients may not have been derived from cancer tissue.

In colorectal cancer, low SAMHD1 levels were detected in tumour tissues relative to adjacent control tissues [Yang et al., 2017], which agrees with a tumour suppressor function. However, higher SAMHD1 levels were associated with metastasis formation [Yang et al., 2017], rather supporting an oncogenic role. The study included the analysis of TCGA data on colorectal cancer [Yang et al., 2017], but no systematic analysis of SAMHD1 across different cancer entities.

The 15 relevant secondary literature articles all had narratives focussing on the potential role of SAMHD1 as a tumour suppressor (Supplementary Table 13).

## Discussion

SAMHD1 has been suggested to exert tumour suppressor functions due its role in maintaining genome integrity and as an inhibitor of uncontrolled proliferation [Herold et al., 2017; Chen et al., 2021]. However, our analysis of TCGA and TARGET data does not suggest that SAMHD1 should be regarded as a *bona fide* tumour suppressor. Notably, SAMHD1 mutations that interfere with SAMHD1 function were not associated with poor outcome, which is something that would be expected from a tumour suppressor. In agreement, no increased cancer formation has been described in SAMHD1-deficient animal models [Kohnken et al., 2015].

Our results rather indicated that changes in SAMHD1 are involved in the oncogenic process in a minority of cases and that it may exert pro-or antitumourigenic effects in different cancer types (and perhaps individual tumours). Moreover, the role of SAMHD1 may differ between the sexes and different races. These findings also show that our understanding of the processes underlying cancer needs to improve further, before a broad paradigm shift towards tumour-agnostic approaches [Danesi et al., 2021] can become a reality.

Notably, the interpretation of our findings may be affected by SAMHD1 being a triphosphohydrolase that cleaves and inactivates the triphosphorylated forms of a number of nucleoside analogues including cytarabine, decitabine, and nelarabine [Schneider et al., 2017; Oellerich et al., 2019; Rothenburger et al., 2020]. However, most cancer diseases are not treated with SAMHD1 substrates. Notably, KIRC (kidney renal clear cell carcinoma), the only cancer in which 5-year survival is significantly lower in SAMHD1 high tumours and SAMHD1 levels are significantly higher in tumour than in control tissues, suggesting an oncogenic role of SAMHD1, is not treated with nucleoside analogues [Geynisman et al., 2021]. Hence, the absence of tumour suppressor activity and/ or oncogenic effects cannot simply be explained by SAMHD1-mediated inactivation of nucleoside analogue substrates.

Our findings demonstrating that SAMHD1 plays multifaceted (and often, if any, minor) roles in cancer seem to be in disaccord with a perception and narrative forming in the field suggesting that SAMHD1 is a tumour suppressor [Herold et al., 2017; Chen et al., 2021]. A systematic review confirmed that most of the available literature focuses on a potential role of SAMHD1 as a tumour suppressor. Among 35 original articles on the role of SAMHD1 in cancer, 31 discussed a potential tumour suppressor function and three potential oncogenic effects. One article reported both potential tumour suppressor and oncogenic activity, but only mentioned the anticipated tumour suppressor effects in the title [Kodigepalli et al., 2018]. All 15 secondary literature articles (reviews, editorials, comments) had a narrative built around SAMHD1 being a candidate tumour suppressor.

The narrative that SAMHD1 is a tumour suppressor has formed since 2013 around findings in a limited number of cancer entities (Supplementary Table 13). Three reasons may contribute to the perpetuation of such a narrative without much scrutiny. Firstly, SAMHD1 has been described to maintain genome integrity by a range of different mechanisms [Herold et al., 2017; Akimova et al., 2021; Chen et al., 2021; Park et al., 2021]. Hence, a potential tumour suppressor role is plausible and convincing. Further research will have to show why SAMHD1’s multifaceted roles in DNA repair do not translate into a consistent and general tumour suppressor function.

The second potential reason is confirmation bias. Scientists (like everybody else) tend to accept findings that support their own experiences, assumptions, and perceptions and to disregard evidence that challenges them [Letrud & Hernes, 2019; Yanai & Lercher, 2021]. Thus, researchers are more likely to look for data that support their hypothesis and not for those that contradict it. Notably, one study referred to SAMHD1 as "novel tumour suppressor" in the title, although *SAMHD1* expression was described as an adverse prognostic factor in Hodgkin lymphoma [Xagoraris et al., 2021].

The final potential reason is publication bias, i.e. a focus on ’positive’ findings that are easier to publish in more prestigious journals than ’negative’ findings [Begley & Ioannidis, 2015; Nissen et al., 2016; Wass et al., 2019; Marks-Anglin & Chen, 2020]. In the case of studies investigating a potential role of SAMHD1 in cancer, this means that some studies that did not find a relationship between SAMHD1 and cancer may simply not have been published and that the publicly available data may not reflect all available data on the subject.

In conclusion, SAMHD1 can play multifaceted roles in cancer that may differ between different cancer types, the sexes, and races. In contradiction to the predominant narrative, SAMHD1 may exert oncogenic as well as tumour suppressor activity and may often be a minor (if any) player in carcinogenesis. Our findings emphasise that hypotheses, perceptions, and assumptions need to be continuously challenged by using all available data and evidence. In this context, it is important that all data are actually published and made available, even if they are not deemed particularly exciting by researchers. Finally, the increasing number of available data and databases should be effectively used to inform and challenge our research and research findings.

## Methods

### Gene Expression and Clinical Data

Gene expression data (FPKM values) from patient tumours were derived from The Cancer Genome Atlas (TCGA) [Cancer Genome Atlas Research Network, 2008] via the GDC Data Portal (https://portal.gdc.cancer.gov). The Bioconductor R package TCGAbiolinks was used to obtain corresponding clinical data. Primary tumour gene expression data and clinical response data were available for 9,572 patients (5,037 female, 4,535 male) with 33 different cancer types. Ages at diagnosis ranged from 14 to 90 (median age at diagnosis = 61, no data for 113 patients). Data were also downloaded for 694 matched normal tissue samples.

Gene expression (RPKM) values and clinical data were extracted for patients in the TARGET database from the National Cancer Institute Office of Cancer Genomics TARGET data matrix (https://ocg.cancer.gov/programs/target/data-matrix). Primary tumour sample data was available for a total of 1,091 patients in TARGET (470 females and 593 males) with seven cancer types. Ages at diagnosis ranged from six days to 32.41 years (median age at diagnosis was 5.4 years (1976 days)).

Tumour vs normal sample gene expression was compared using the wilcox.test function in R, which performs the Mann Whitney U test for independent groups. Pairwise comparisons were made using the Wilcoxon Signed Rank test. Pie charts were generated using ggplot2.

### Methylation and miRNA data

TCGA methylation beta values and miRNA expression values (reads per million miRNA mapped) were downloaded from the GDC Data Portal (https://portal.gdc.cancer.gov). Mean methylation beta values for each CpG site in the SAMHD1 promoter for which data were available (cg02078758, cg00642209, cg16430572, cg09128050, cg12099051, cg18861300, cg11094122, cg22769031, cg23888977, cg09717261, cg24951864, cg06097592, cg22583967, cg10804363 and cg12517061) were calculated per individual. Expression data for miRNAs which were listed in DIANA-TarBase v8 [Karagkouni et al., 2018] as being experimentally validated to positively interact with SAMHD1 (n=21) along with eight miRNAs shown in previous experiments to target SAMHD1 [Jin et al., 2014; Pilakka-Kanthikeel et al., 2015; Kohnken et al., 2017; Riess et al., 2017] were extracted for analysis. Scatter plots and associated Pearson correlations for methylation and miRNA expression with SAMHD1 expression were calculated using the ggplot2 package in R.

### Survival analyses

Cox proportional hazards regression was used to calculate the hazard ratio for cohorts expressing high levels of SAMHD1. Overall survival (OS) was defined as days to last follow-up or death, as previously described [Ng et al., 2016]. Calculations were performed using the R survminer and survival packages. The ‘surv_cutpoint’ function was used to identify the optimal expression cut-off point to give the lowest p-value for high vs low expression. We permitted the cut-off to be only between the 20^th^ and 80^th^ percentiles of gene expression values, as described by previously [Uhlen et al., 2017].

Kaplan-Meier survival curves were generated using R package ggsurvplot. P-values in each case were the result of a log rank (Mantel-Cox) test, which assesses whether there is a significant difference between the survival of two independent groups. Hazard ratios quoted refer to values for ‘low’ (below the calculated optimal cut-off) expression for each gene in the model, with values >1 indicating increased hazard (i.e. reduced OS) and values <1 indicating decreased hazard (i.e. increased OS).

### Mutation Data and Variant Effect Prediction

Mutation data for 10,149 TCGA patients were downloaded from the GDC Data Portal (https://portal.gdc.cancer.gov). A dot plot displaying mutation frequencies of 21,156 genes was generated using ggplot2.

In order to assess the potential impact of mutations in SAMHD1, we used the online tool Variant Effect Predictor (VEP) [McLaren et al., 2016] to obtain reports from SIFT [Sim et al., 2012], PolyPhen-2 [Adzhubei et al., 2010], Condel [González-Pérez & López-Bigas, 2011] and CADD [Kircher et al., 2014].

A lollipop plot of SAMHD1 mutations was generated using the cBioPortal MutationMapper tool (https://www.cbioportal.org/mutation_mapper) [Cerami et al., 2012].

### Literature review

Relevant articles were identified on 17^th^ June 2021 by using the search term "(((Cancer) OR (tumor) OR (tumour))) AND (SAMHD1)" in PubMed (https://pubmed.ncbi.nlm.nih.gov) on the basis of the principles outlined in the PRISMA guidelines (http://prisma-statement.org). Articles in English were included into the analysis, when they contained original data on the role of SAMHD1 in cancer. Moreover, reviews that discussed the potential impact of SAMHD1 on cancer were used to analyse conceptions and the predominant narrative in the field. Two reviewers independently analysed articles for relevant information and then agreed a list of relevant articles.

### Conflict of interest

There is no conflict of interest.

## Notes

### Competing Interest Statement

The authors have declared no competing interest.

### Summary of Updates

Correct some minor errors, such as typos.

