## Supplementary material for "Multifaceted roles of SAMHD1 in cancer": Suppl Tables and Figures: McLaughlin et al_Suppl Figure 1.pdf

### Supplementary Figure 1

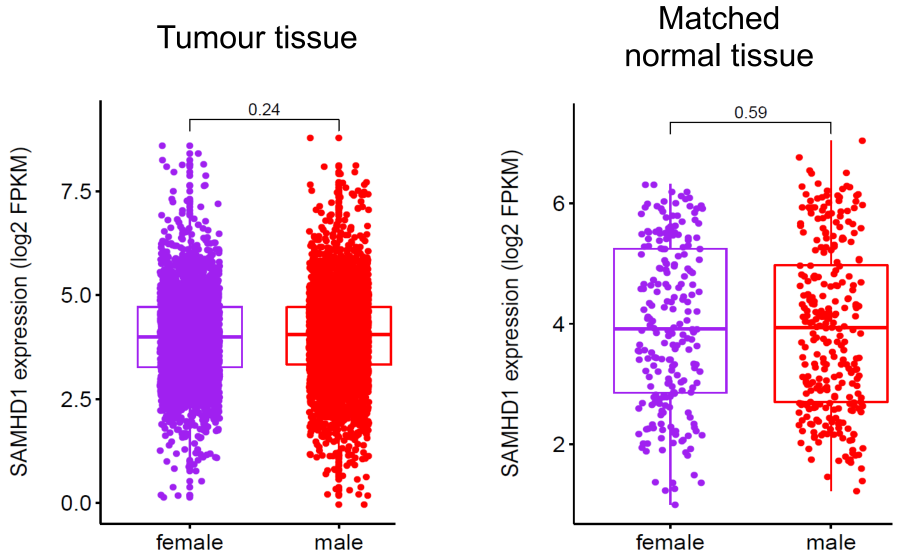


**Supplementary Figure 1**. Comparison of *SAMHD1* expression levels as expressed by fragments per kilobase of transcript per million mapped reads (FPKM) after removal of sex-specific cancer types and of BRCA (breast invasive carcinoma), for which only a very small fraction of samples (12/ 1,089 tumour tissue samples, 1/ 113 matched normal tissue samples) was derived from males. P-values were determined by Mann-Whitney U (Wilcoxon) test for independent groups.
