## Supplementary material for "Multifaceted roles of SAMHD1 in cancer": Suppl Tables and Figures: McLaughlin et al_Suppl Figure 2.pdf

### Supplementary Figure 2

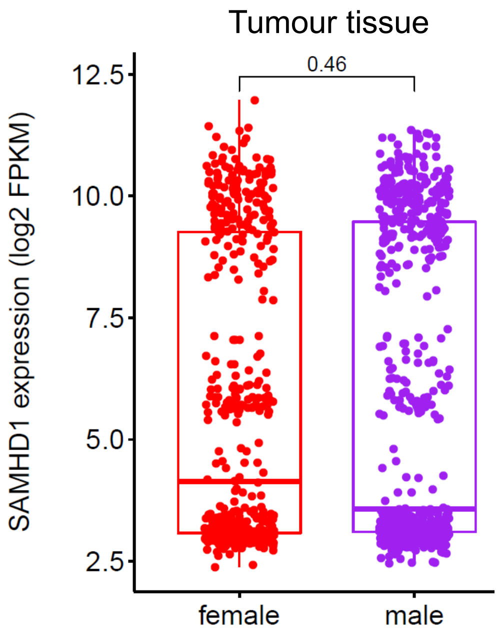


**Supplementary Figure 2**. Comparison of *SAMHD1* expression levels as expressed by fragments per kilobase of transcript per million mapped reads (FPKM) in tumours from the TARGET database. P-value determined by Mann-Whitney U (Wilcoxon) test for independent groups.
