## Supplementary material for "Multifaceted roles of SAMHD1 in cancer": Suppl Tables and Figures: McLaughlin et al_Suppl Figure 3.pdf

### Supplementary Figure 3

A

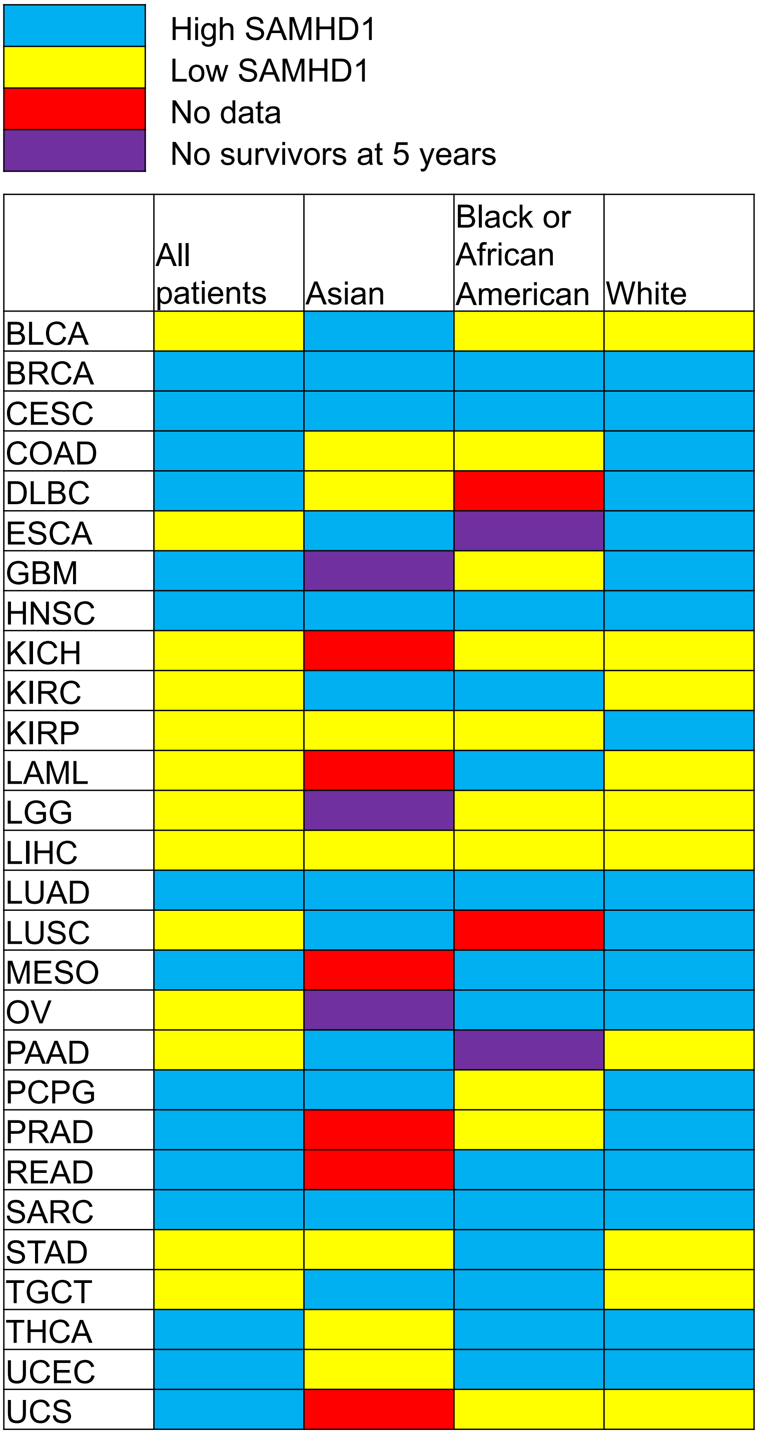


**Suppl. Figure 3A**. Heatmap indicating the association of SAMHD1 expression and 5-year survival rates (blue: high SAMHD1 associated with higher survival rates, yellow: low SAMHD1 associated with higher survival rates).

### Supplementary Figure 3

B

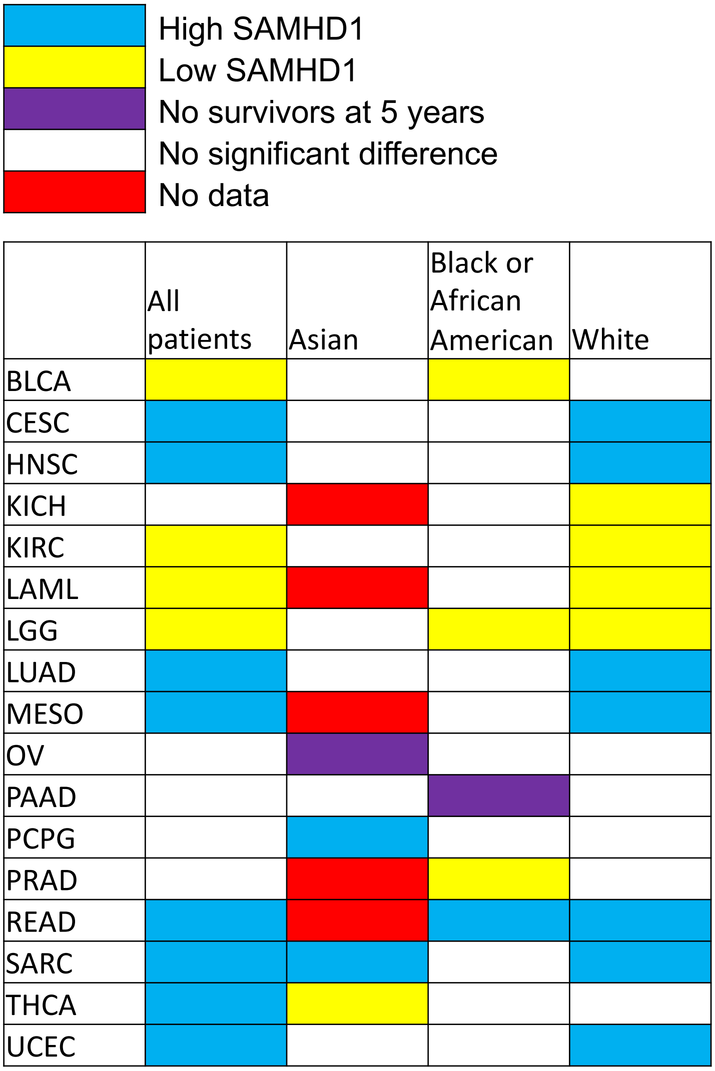


**Suppl. Figure 3B**. Heatmap indicating cancer entities in which high SAMHD1 expression (blue) or low SAMHD1 expression (yellow) is significantly ($p<0.05$) associated with higher 5-year survival rates.
