## Supplementary material for "Multifaceted roles of SAMHD1 in cancer": Suppl Tables and Figures: McLaughlin et al_Suppl Figure 5.pdf

### Supplementary Figure 5

A

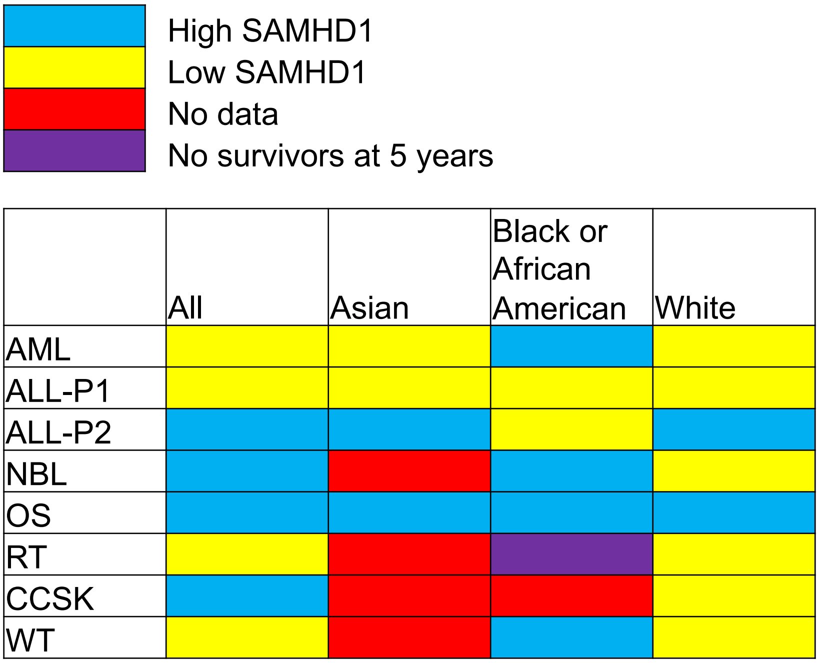


B

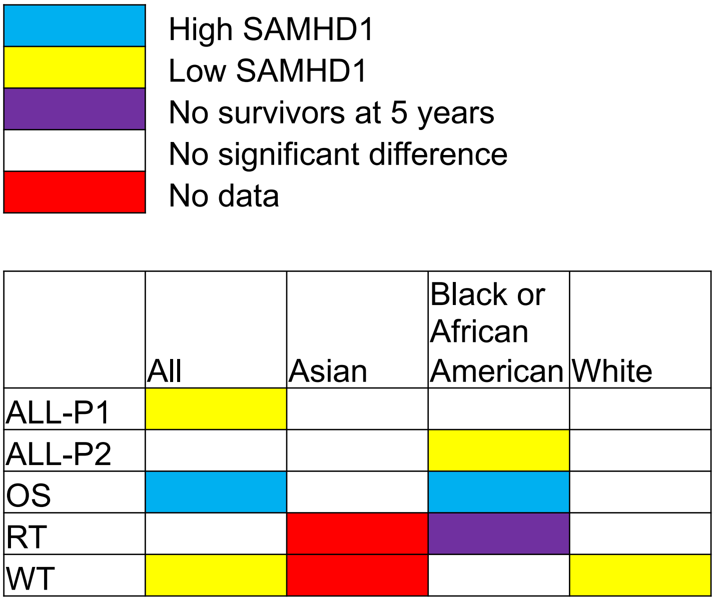


**Suppl. Figure 3**. Heatmap indicating the association of SAMHD1 expression and 5-year survival rates (blue: high SAMHD1 associated with higher survival rates, yellow: low SAMHD1 associated with higher survival rates). A) All cancer types independently of significance level. B) Cancer types in which at least comparison resulted in a significant ($p<0.05$) difference.
