## Supplementary material for "Multifaceted roles of SAMHD1 in cancer": Suppl Tables and Figures: McLaughlin et al_Suppl Figure 6.pdf

### Supplementary Figure 6

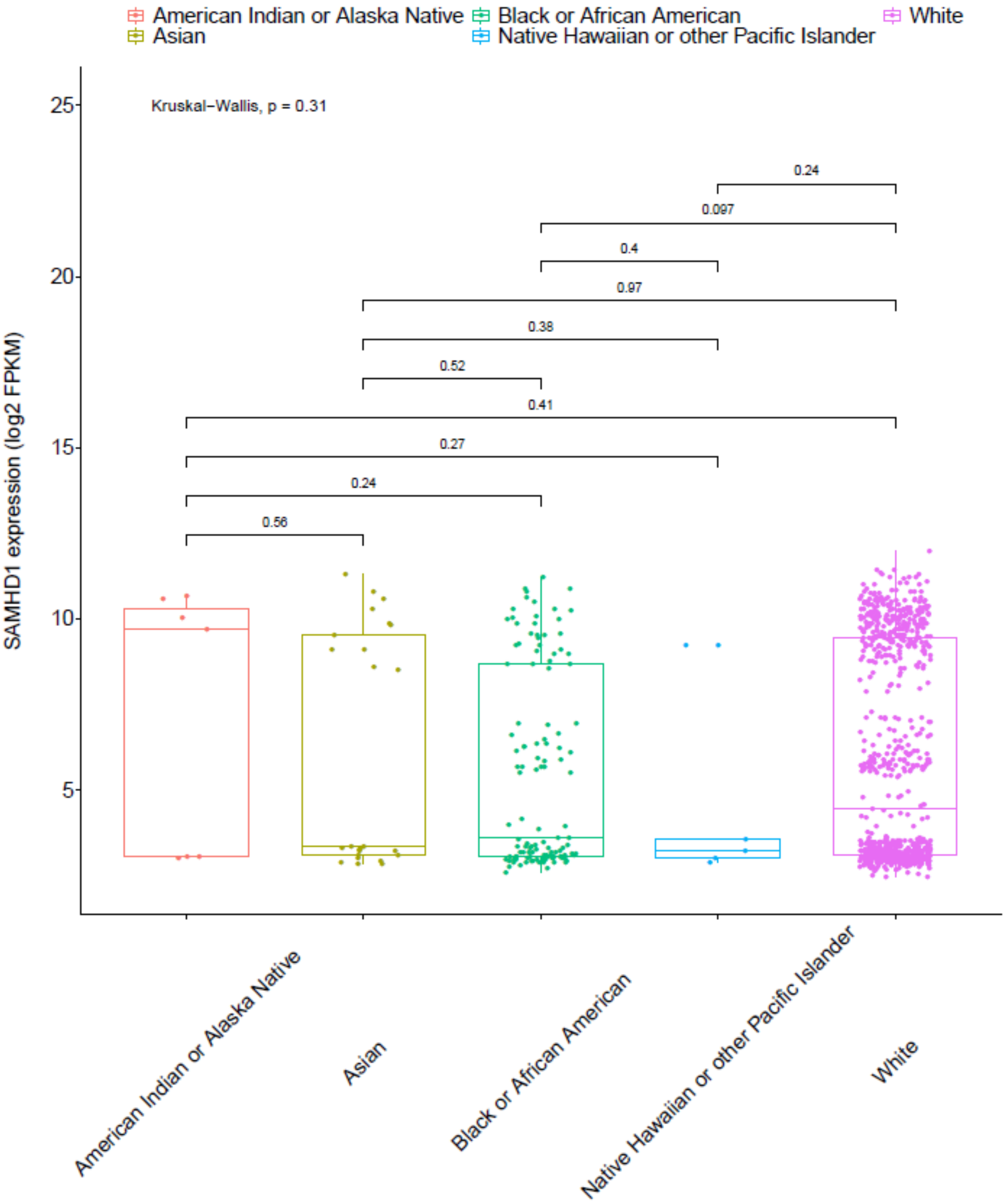


**Supplementary Figure 6**. SAMHD1 levels in patients of different race in the TARGET database. Comparison of all groups was performed using the Kruskal-Wallis test. Individual group comparisons were performed using the Mann-Whitney U (Wilcoxon) rank sum test.
