## Supplementary material for "Multifaceted roles of SAMHD1 in cancer": Suppl Tables and Figures: McLaughlin et al_Suppl Figure 7.pdf

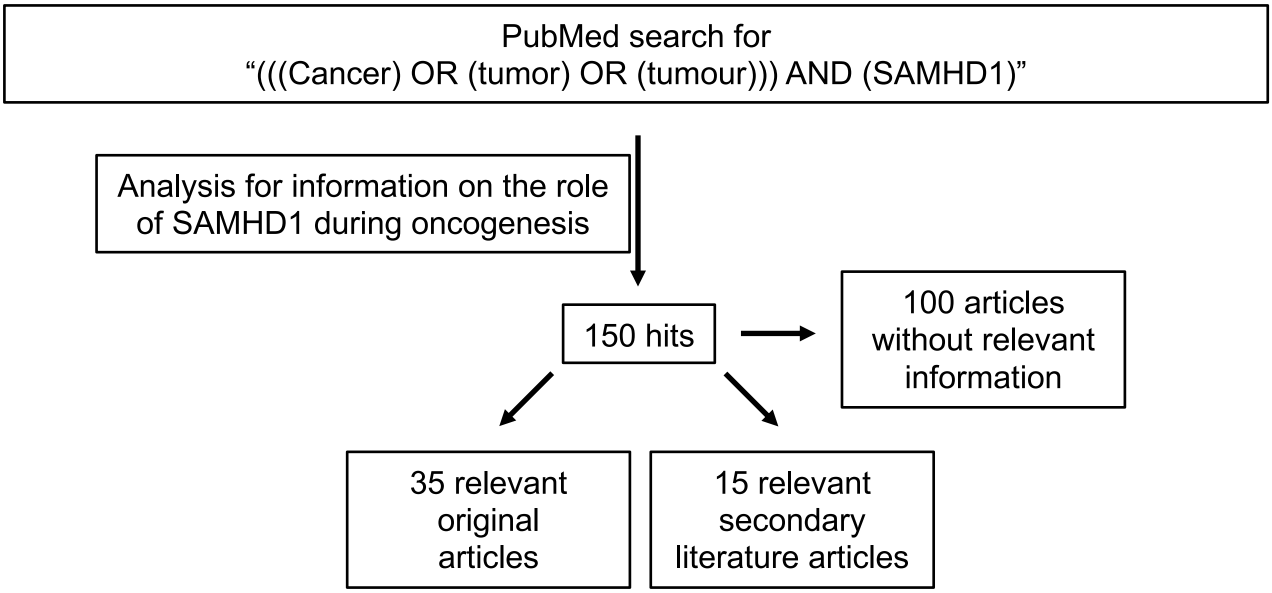


**Supplementary Figure 7**. Summary of the findings of the literature review for articles containing data on the role of SAMHD1 during oncogenesis. Articles were identified by using the search term "(((Cancer) OR (tumor) OR (tumour))) AND (SAMHD1)" in PubMed (https://pubmed.ncbi.nlm.nih.gov) on 17th June 2021.
